# A dense linkage map for a large repetitive genome: discovery of the sex-determining region in hybridising fire-bellied toads (*Bombina bombina* and *B. variegata*)

**DOI:** 10.1101/2020.10.06.328633

**Authors:** Beate Nürnberger, Stuart J.E. Baird, Dagmar Čížková, Anna Bryjová, Austin B. Mudd, Mark L. Blaxter, Jacek M. Szymura

## Abstract

Hybrid zones that result from secondary contact between diverged populations offer unparalleled insight into the genetic architecture of emerging reproductive barriers and so shed light on the process of speciation. Natural selection and recombination jointly determine their dynamics, leading to a range of outcomes from finely fragmented mixtures of the parental genomes that facilitate introgression to a situation where strong selection against recombinants retains large unrecombined genomic blocks that act as strong barriers to gene flow. In the hybrid zone between the fire-bellied toads *Bombina bombina* and *B. variegata* (Anura: Bombinatoridae), two anciently diverged and ecologically distinct taxa meet and produce abundant, fertile hybrids. The dense linkage map presented here enables genomic analysis of the selection-recombination balance that keeps the two gene pools from merging into one. We mapped 4,775 newly developed marker loci from bait-enriched genomic libraries in F2 crosses. The enrichment targets were selected from a draft assembly of the *B. variegata* genome, after filtering highly repetitive sequences. We developed a novel approach to infer the most likely diplotype per sample and locus from the raw read mapping data, which is robust to over-merging and obviates arbitrary filtering thresholds. Large-scale synteny between *Bombina* and *Xenopus tropicalis* supports the resulting linkage map. By assessing the sex of late-stage F2 tadpoles from histological sections, we also identified the sex-determining region in the *Bombina* genome to 7 cM on LG5, which is homologous to *X. tropicalis* chromosome 5, and inferred male heterogamety, suggestive of an XY sex determination mechanism. Interestingly, chromosome 5 has been repeatedly recruited as a sex chromosome in anurans with XY sex determination.

## Introduction

When two genetically differentiated populations come into contact and produce fertile hybrids, any existing reproductive barriers between them are tested. Theory predicts that unless these barriers are strong and based on many loci across the genome, recombination in the newly formed hybrid zone will with time break up ancestral haplotypes into ever smaller segments (Barton 1983; Barton and Bengtsson 1986). As a result, variants at neutral loci will become dissociated from loci whose alleles are barred by natural selection from introgressing into the opposite gene pool. While neutral variation is eventually eroded, stable allele frequency clines should remain at loci under selection. But this separation of fates is a slow process that may take thousands of generations to complete (Baird 1995; Kruuk et al. 1999). Prior to that, the width and shape of clines, even at neutral markers, can inform about the balance between gene flow, selection, and recombination in a given hybrid zone (Barton and Gale 1993). An even more detailed picture emerges from the length distribution of local ancestry tracts, *i.e*. haplotype segments inherited from one taxon and bounded by recombination breakpoints. For a given distribution, likely combinations of hybrid zone age and selection regime may be inferred (Baird 1995). Local, transient distortions in the length distribution may pinpoint genomic regions under strong selection (Sedghifar et al. 2016). Local ancestry tracts are the natural units of inheritance in a hybrid zone (Baird 2006) and can be inferred from dense linkage maps. To access this rich source of information, we developed a linkage map from F2 crosses of the fire-bellied toads *Bombina bombina* and *Bombina variegata*, a textbook example (Urry et al. 2020) of hybridisation between two anciently diverged taxa.

Local ancestry tracts provide the most direct evidence for hybridisation, as they cannot be explained by incomplete lineage sorting or convergence (Rieseberg et al. 2000). They have been used to infer the number of generations required to stabilise a hybrid sunflower species (Ungerer et al. 1998), uncover the lack of F2 or deeper hybrid generations in a *Populus* hybrid zone (Christe et al. 2016), compare the age of two separate hybrid zones of *Lissotriton* newts (Zieliński et al. 2019), detect past episodes of hybridisation (Meier et al. 2017; Węcek et al. 2017; Duranton et al. 2020), localise introgressed genomic segments (Huerta-Sánchez et al. 2014; vonHoldt et al. 2016), identify incompatible haplotype combinations in hybrid swordfish (Powell et al. 2020), and monitor shifts in genome composition in experimental *Drosophila* populations (Matute et al. 2020). The rapidly growing theoretical literature infers evolutionary processes from genome-wide local ancestry patterns, from the age of ancient admixture pulses (Harris and Nielsen 2013) to the onset of neutral mixing with continuous gene flow (Sedghifar et al. 2015), adaptive introgression (Sachdeva and Barton 2018; Shchur et al. 2019), and selection against deleterious allele combinations in hybrid zones (Sedghifar et al. 2016; Hvala et al. 2018).

The fire-bellied toad *B. bombina* and the yellow-bellied toad *B. variegata* hybridise in typically narrow (2 – 7 km wide) contact zones wherever their ranges adjoin in Central and Eastern Europe (Yanchukov et al. 2006). Transcriptome-based coalescence analyses suggest that their lineages split no later than 3.2 million years ago (Ma) (Pabijan et al. 2013; Nürnberger et al. 2016). They profoundly differ in a large number of traits, many of which are likely adaptations to different habitats (Szymura 1993): *B. bombina* reproduces in semi-permanent lowland ponds, whereas *B. variegata* is adapted to ephemeral aquatic sites, typically at higher elevations. The hybrid zones are maintained by natural selection and pose barriers to neutral gene flow, as evidenced by a sharp central allele frequency step in geographic clines, strong linkage disequilibria between independently segregating genetic markers, and cline stability over 50 and 70 year sampling intervals (Szymura and Barton 1991; Yanchukov et al. 2006). From cline shape, Szymura and Barton (1991) estimated that central hybrid populations had a 42% lower fitness than the pure taxa, consistent with incompatibilities at dozens of loci. Under uniform experimental conditions, embryo and tadpole survival is lower in hybrids than in the pure taxa (Kruuk et al. 1999b). Despite these fitness effects, detailed analyses of transects in Poland, Croatia, Romania and Ukraine based on a small (< 10) number of loci uncovered a wide range of recombinants, with F1s nearly if not entirely absent (see Yanchukov et al. 2006 for a summary). We wish to explore this mosaic of ancestry blocks within individuals and across the hybrid zone to better understand the conundrum of abundant hybridisation despite ancient divergence.

Based on current technology, *Bombina*’s large and repetitive genome (7-10 Gb, Gregory 2020) precludes population genomic analysis using whole genome sequencing and hampered a previous attempt to generate a linkage map (Nürnberger et al. 2003). We therefore opted for targeted enrichment (reviewed in Jones and Good 2016) based on a new draft assembly of a *B. variegata* genome and published *Bombina* transcriptomes (Nürnberger et al. 2016), and we applied this to a controlled, three-generation experimental cross between *B. variegata* and *B. bombina*. This reduced representation approach (Davey et al. 2011) allowed us to filter out repetitive regions before selecting enrichment targets, obviated the need to infer exon-intron boundaries (as in exome capture, Neves et al. 2013) and, compared to methods based on restriction enzyme digests, promised greater reproducibility and more even target coverage for this large genome (Jones and Good 2016). *Bombina* belongs to the superfamily Discoglossoidea, which split ~200 Ma from other anuran lineages with available genome assemblies (Feng et al. 2017). Capture probes derived from *Xenopus* or *Hyla* are thus not expected to work well in *Bombina* (Hedtke et al. 2013; Hutter et al. 2019). Enrichment success across taxon boundaries declines sharply in the range of 5-10% absolute sequence divergence, d_xy_ (Hedtke et al. 2013; Jones and Good 2016; Hutter et al. 2019). The distribution of d_xy_ between *B. bombina* and *B. variegata* has a mean of 0.0202 and a mode at 0.013 (Nürnberger et al. 2016). We therefore expect reliable cross-taxon enrichment for the great majority of targets as well as an abundant supply of ancestry-informative markers.

Read coverage of a given enrichment target is typically highest in the centre and drops off at the ends (Chevalier et al. 2014; Harvey et al. 2016), and thus variants can vary widely in their read support. Moreover, erroneously mapped reads can produce spurious signal of variation (McCartney◻Melstad et al. 2016). Different variants, when called separately, can therefore produce contradictory signals for the same target and sample. Instead of censoring data by setting arbitrary filtering thresholds, we use the total information contained in reads mapped to a given target and, for each sample, computed the likelihood of three possible diplotypes: *B. bombina* homozygote (BbHOM), heterozygote (HET), and *B. variegata* homozygote (BvHOM). To this end, we polarised the raw read mapping data so as to maximise the difference between the grandparents, a *B. variegata* male and a *B. bombina* female. Across all reference positions of a given target, sequence states associated more with one grandparent than the other were weighted by their read support and contribute to separate scores of ‘*bombina*-ness’ and ‘*variegata*-ness’, respectively. When these scores are plotted in a coordinate system, samples cluster by diplotype, with homozygotes near x and y axes and heterozygotes along or near the diagonal. Using this clustering and an explicit genetic model, we inferred the most likely diplotypes and propagated their statistical support to the map-making stage.

We coupled the new linkage map with further data to answer two questions. First, we analysed the homology of the molecular bait sequences against the *Xenopus tropicalis* genome. The large-scale synteny across ~220 million years of anuran evolution describes aspects of the likely Bombinanura ancestral chromosome state and serves as a quality check of the map. Second, we coupled diplotype estimates with histological estimates of F2 progeny sex; sex-biased segregation allowed us to locate the sex-determining (SD) on the *Bombina* map and infer the SD mechanism. As is true for 96% of amphibians (Eggert 2004), *Bombina* lacks heteromorphic sex chromosomes. Frequent turnover of sex chromosomes (Miura 2017; Jeffries et al. 2018) and/or very rare X-Y (or Z-W) recombination events, *e.g.* in sex-reversed females, (Perrin 2009; Stöck et al. 2011; Guerrero et al. 2012; Rodrigues et al. 2018) may counteract the expected degeneration of the Y (or W) chromosome (Charlesworth and Charlesworth 2000) in this clade. Biased hybrid sex ratios are thought to have prompted the establishment of two new SD systems, one with male heterogamety and the other with female heterogamety, in the Japanese wrinkled frog *Glandirana rugosa (Miura 2017)*. Given the strong selection on and rapid divergence of SD systems (Coyne and Orr 2004), the map location of the *Bombina* SD region will be important for our analyses. In some hybrid zones, sex-linked as opposed to autosomal loci have formed steeper clines suggestive of stronger gene flow barriers (Oryctogalus, Carneiro et al. 2013; Gryllus, Maroja et al. 2015; Hyla. Dufresnes et al. 2016). On the other hand, striking cases of sex-linked introgression have been found and attributed to genetic conflict over the sex ratio (Mus, Macholán et al. 2008; Drosophila, Meiklejohn et al. 2018). Knowledge of the location of the SD region in *Bombina* will thus be critical for the analysis of the hybrid zone.

## Materials and Methods

### Laboratory crosses

A male *B. v. variegata* from Obidowa (near Nowy Targ, Poland, sample acc. # ERS3926742) was crossed with a female *B. bombina* from Wodzisław Małopolski (Poland, sample acc. # ERS3926743) in 2014. Eighty F1 offspring were raised to maturity, and one F1 male was crossed with two F1 females to produce two F2 families (families 6 and 7 in the following, see File S1 for husbandry, offspring rearing and F1 sample accessions). The F2 offspring were raised to advanced metamorphosis (Gosner stages 42-44, Gosner 1960) and were humanely killed by MS222 (Ethyl 3-aminobenzoate methanesulfonate) overdose. For 80 offspring of family 6 and 82 offspring of family 7, the gonads with mesonephroi were dissected and fixed in Bouin’s solution (Kiernan 1990), while the remaining tissue was frozen. Toe clips were collected from the *B. bombina* grandmother and each of the F1 offspring under MS222 anesthesia. The *B. variegata* grandfather was euthanised by MS222 overdose and dissected for whole genome sequencing. Tissue samples for DNA extraction were kept at −80 °C.

### Whole genome sequencing

DNA was extracted from muscle tissue of the *B. variegata* grandfather using the Invisorb Spin Tissue Minikit (Stratec, Germany). PCR-free TruSeq libraries with mean insert sizes of 350 bp (n = 8) and 550 bp (n = 2) were prepared by Edinburgh Genomics and sequenced on the Illumina HiSeq X, producing 6.67 × 10^9^ (350 bp) and 1.05 × 10^9^ (550 bp) read pairs (150 bp, PE). Adapter removal and quality trimming were carried out with bbduk (BBMap suite v.36.76, B. Bushnell, sourceforge.net/projects/bbmap/). Parameters for adapter removal were k=23, mink=8, and edist=1 for R1 and k=23, mink=8, and edist=2 for R2. Quality trimming parameters were trimq=20, maq=25, and minlength=50. Genome size was estimated from unassembled reads with the preqc module of the String Graph Assembler (SGA, v. 0.10.15) (Simpson and Durbin 2012; Simpson 2014) using a subset of 1.1 × 10^9^ read pairs. All libraries were evenly represented in this and subsequent subsets

### Genome Assemblies

A subset of 1.29 × 10^9^ read pairs (approximately 45× genome coverage) were assembled with the CLC Genomics Workbench (v. 9.5.3) (Qiagen, Hilden, Germany) using default parameters. Repeat sequences were assembled with REPdenovo (v. 2017-02-23) (Chu et al. 2016) with default parameters except MIN_REPEAT_FREQ=100 (Chong Chu, pers. comm.). REPdenovo produced an unmerged version of all assembled repeats and a merged version by combining repeats with more than 90% identity. All quality-trimmed reads were mapped to the unmerged REPdenovo output with Bowtie2 (v. 2.2.3) (Langmead and Salzberg 2012), and the 52% of read pairs that did not map were extracted as the repeat-subtracted read set. We queried the merged REPdenovo output against Repbase (Jurka et al. 2005; Bao et al. 2015) with the Censor tool (Kohany et al. 2006, blastn and tblastx, vertebrate database, last accessed 31 July 2020). Following Rogers et al. (2018), we annotated each merged REPdenovo contig with the highest scoring match and mapped a subset of 7.43 × 10^7^ read pairs (approximately 2.64× genome coverage) to the merged REPdenovo output with Bowtie2 (v. 2.2.3) (Langmead and Salzberg 2012). Mean mapped read coverage was divided by 2.64 to estimate copy number.

The repeat-subtracted read set was assembled with SGA and Platanus, and sequences identical in these new assemblies and the previous CLC assembly were considered for bait design. For the SGA (v. 0.10.15) (Simpson and Durbin 2012) assembly, we followed the steps in the example assembly of a human genome (see the ../src/examples/ directory of the SGA distribution) using a subset of 1.12 × 10^9^ read pairs (approximately 40× genome coverage). For the Platanus (v. 1.2.4) (Kajitani et al. 2014) assembly, we extracted CLC contigs that matched the published *B. v. variegata* transcriptome (Nürnberger et al. 2016) and 125 gene sequences from public databases based on a minimum sequence identity of 90% with BLAST+ (v. 2.2.3) (Camacho et al. 2009). Reads that mapped to the extracted CLC contigs with Bowtie2 (v. 2.2.3) (Langmead and Salzberg 2012) were assembled with the Platanus (v. 1.2.4) (Kajitani et al. 2014) assemble step.

### Candidate sequences and bait design

Candidate sequences for bait design were selected from the CLC assembly based on uniqueness, correct assembly, and minimal redundancy. We considered subsets of CLC contigs to be unique if they did not have any matches to other CLC contigs, based on an 85% sequence identity threshold with BLAST+ (v. 2.2.3) (Camacho et al. 2009). CLC contig sequences with exact matches (minimum length 100 bp) in the SGA and Platanus assemblies were deemed correctly assembled. Coverage and variant information (‘bubbles’) provided by Platanus was used to flag overmerged sequences (see File S1 for details). To minimise the proximity of enrichment targets (local redundancy), the CLC assembly was scaffolded against the *B. v. variegata* transcriptome assembly (Nürnberger et al. 2016) using SCUBAT2 (G. Koutsovoulos, https://github.com/GDKO/SCUBAT2, commit b03e770). For each SCUBAT2 path (*i.e*. a set of contigs linked by exons from a single transcript), we identified the longest sequence section that was unique, correct, and lacked excessive variation. We also selected candidate sequences in CLC contigs (minimum length 5 kb) that were not included in any SCUBAT2 paths. These were filtered as previously described, except that exact matches were not confirmed against the Platanus assembly. Finally, all candidate sequence positions with a BLAST+ (v. 2.2.3) (Camacho et al. 2009) alignment against the unmerged REPdenovo output were hard masked.

We submitted 6,400 candidate sequences (minimum length 500 bp; 4,400 with known gene association) to Arbor Biosciences (Ann Arbor, Michigan, USA) for bait design and synthesis. For each of 5,000 enrichment targets, four 100 base baits were designed that aligned with 50 base offsets to a 250 base sequence stretch (2x tiling). Baits were designed according to the strictest in-house criteria (no BLAST+ match to the CLC assembly with T_m_ > 60° C, no ‘N’ positions, %GC between 25 and 55, no RepeatMasker matches, and ΔG > −8).

### Enriched genomic libraries and sequencing

Genomic DNA was extracted from the F0 *B. bombina* grandmother, the three F1 parents, and the 162 F2 offspring using the Invisorb Spin Tissue Minikit (Stratec, Germany). DNA concentrations were measured by Qubit fluorometer (Invitrogen, USA) and normalized to 50 ng/μl. DNA extractions were then fragmented with the Bioruptor Pico (Diagenode, Belgium) using 7 cycles of 30s fragmentation and 60s cooling, which resulted in a mean fragment length of approximately 250 bp. Libraries were constructed from the fragmented DNA using the KAPA HyperPrep Kit (Kapa Biosystems, South Africa) per the manufacturer’s instructions, except all reaction volumes were halved. Dual indexed TruSeq-like adapters were added by ligation of “universal stubs”, followed by 8 cycles of PCR using indexed primers, as described by (Glenn et al. 2019). SpriSelect beads (Beckman Coulter, USA) were used to size select the libraries, eliminating high molecular weight fragments with a 0.6x bead to sample volume ratio and low molecular weight fragments with a 1x ratio. Libraries were pooled in equimolar ratios (number of samples: 1, 2, or 4) and concentrated to 7 μl with 1x SpriSelect beads. The library pools were enriched using the myBaits target capture kit (Arbor Biosciences, Ann Arbor, Michigan, USA) with the custom baits. Hybridisation was run at 65°C for 20 hr. Enriched libraries were amplified with universal P5 and P7 primers during 11 cycles of PCR (PCR conditions as per the KAPA HyperPrep Kit). Amplified libraries were purified using 1x SpriSelect beads and mixed in equimolar ratios.

We tested the enrichment success and the effect of pooling libraries (1, 2, or 4 per enrichment reaction, including mixtures of the two taxa) using a single run of the Illumina MiSeq (v2 flow cell, 150 bp, PE). Because there was no apparent detriment to enriching four libraries in one reaction, this level of pooling was used for the entire dataset, excluding four instances with fewer than four samples. Enriched libraries of the *B. bombina* grandmother, the three F1 parents, and all 162 F2 offspring were sequenced on one lane of the Illumina NovaSeq (S1 flow cell, 150 bp, PE) by Edinburgh Genomics. An enriched library of the *B. variegata* grandfather was included in the Miseq test.

### Mapping reference

Because the enriched libraries span beyond the 250 bp bait regions, we used the ‘Assembly by Reduced Complexity’ (ARC) package (v. 1.1.4-beta) (Hunter et al. 2015) to determine the mapping reference for each target. ARC bins read pairs based on the bait region to which they map and computes a unique *de novo* assembly for each bin with SPAdes (v. 3.9.0) (Nurk et al. 2013). This process is iterative, with the last *de novo* assembly used as the reference for the next mapping round until contig lengths stop increasing. From the enriched read-set of the F0 *B. variegata* adult, assemblies were obtained for 4,850 targets. These were aligned against the CLC target contigs using BLAST+ (v. 2.2.3) (Camacho et al. 2009) in order to eliminate any sequence erroneously added to assembly termini and to resolve chimeric assemblies (McCartney◻Melstad et al. 2016). This screen resulted in mapping references for 4,763 targets (see File S1 for details). For the remaining 237 targets, the entire CLC contig was used as the reference. We constructed an analogous mapping reference for the *B. bombina* grandmother.

### Read mapping and diplotyping

The enriched sequence data were processed as previously described to produce repeat-subtracted reads sets. These were mapped with Bowtie2 (v. 2.2.3) (Langmead and Salzberg 2012) to the *B. variegata* reference and, for a few samples, to the *B. bombina* analogue to estimate mapping bias. Duplicates were flagged with Picard (v. 2.6.0) (Broad Insitute 2019) MarkDuplicates. For each bait interval, the mapped read data was summarised using Samtools (v. 1.4) (Li et al. 2009) mpileup and PoPoolation2 (Kofler et al. 2011) mpileup2sync. The resulting summary files contain, for each sample and locus, a matrix of *n* columns (*n* = number of reference positions) and six rows (sequence states of A, C, G, T, DEL, and N; Figure 1A, B) of the counts of reads supporting each sequence state at each position. Note that insertions cannot be represented in this matrix of *reference* coverage. These summaries were analysed using a “Fast Vector” (FastVec) Mathematica (v. 12.0) (Wolfram Research, Inc. 2019) script: it avoids the computational load of per-reference-position-state estimation combinatorics and positions summary matrices on a linear *bombina-variegata* vector (see file S1 for details). An open source Python version is under development. Briefly, the vector endpoints are calculated in two steps. First, for the two F0 grandparents, the counts are divided by the column totals to obtain frequencies. Subtracting the resulting *B. bombina* frequency matrix from the *B. variegata* frequency matrix gives a polarised matrix where positive entries represent sequence states that are more common in *B. variegata*, and negative entries are states more common in *B. bombina*. Signed entries are then weighted with respect to the support for this distinction in each matrix column (at each position): For a given position *i*, we computed the significance *Sig*(*i*) of the likelihood ratio test on the raw read counts of the two grandparents, comparing the hypotheses they were drawn either from the same or from different multinomial distribution(s). All matrix elements in column *i* were then multiplied by (1 – *Sig(i*)). This gave the initial weighted polarised matrix, ***M***_p_ (Figure 1C). The raw read count matrix for each sample was multiplied by ***M***_p_. The means of the positive and negative entries express the average weighted read coverage of sequence states associated with the *B. variegata* grandfather and the *B. bombina* grandmother, respectively, for that sample. When these positive and negative scores are plotted in a coordinate system, samples at a given locus typically fall into three clusters representing the three diplotypes (BbHOM, HET, and BvHOM; Figure 1D), with low coverage (and/or low power) individuals’ data near the origin.

**Figure 1.**
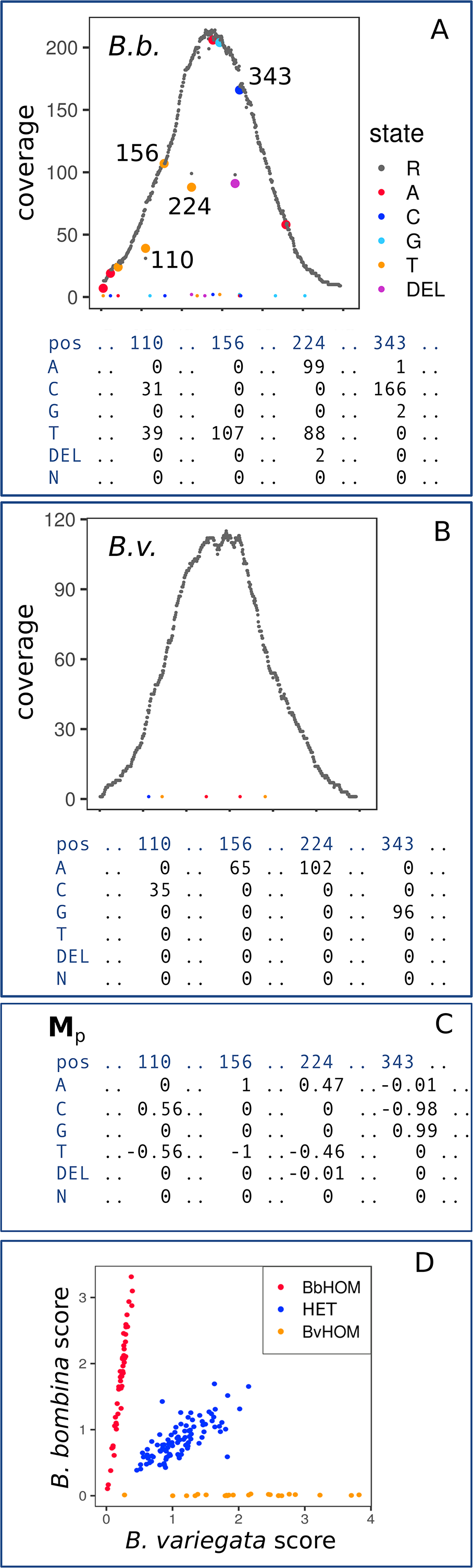
Polarisation of the raw read coverage. Plots show the raw read coverage along the reference sequence (x-axis) of locus 332,172 for F0 *B. bombina* (A) and F0 *B. variegata* (B). The homozygous *B. variegata* diplotype is identical to the locus sequence for this individual (reference state (R) only). Four variant positions (110, 156, 224 and 343) are highlighted, and the raw read counts of the six possible sequence states are noted in the matrices below the plots. A polarised matrix, **M**_p_, is computed from these read counts in two steps (see text, C), in which sequence states associated with *B. variegata* have positive entries and sequence states associated with *B. bombina* have negative entries. For each sample, raw read counts are then multiplied by **M**_p_. Average positive entries and average negative entries result in a *B. bombina* score and a *B. variegata* score, respectively, and when plotted in a coordinate system (D), samples can be assigned to three clusters representing BbHOM, HET, and BvHOM. Note that the heterozygous variants (panel A) do not interfere with the clustering into three diplotypes.

Assuming that the clusters closest to the axes (Figure 1D) represent homozygous diplotypes, the vector endpoints (currently estimated from a single individual each) can be re-estimated from the combined raw count matrices over each of these clusters in a second **M**_p_ estimation step (now based on higher coverage). After this **M**_p_ update, separation of clusters such as (Figure 1D) is unchanged or improved. We reduced the combined read counts of each of these clusters to a strict majority consensus, giving us a set of candidate haplotypes. Truly HOM clusters should result in well supported (high coverage depth) haplotype estimates. For each sample’s read counts, we then computed the parental likelihoods of all possible candidate haplotype combinations, accounting for error, contamination (homozygote clusters: deviation from the 0° and 90°, respectively), and enrichment bias (heterozygote clusters: deviation from 45°). The maximum likelihood candidate haplotype pair (MLCHP) is assumed to be that with the largest total parental likelihood over all individuals. The maximum likelihood diplotype for an individual is reported along with its support estimates with respect to the MLCHP and across all parental candidates (see File S1 for a full description).

We re-scored the 327 (6.5% of the total) loci that did not show the expected diploptypes in the F0 (BvHOM and BbHOM) and F1 (HET, HET, and HET) individuals. For each locus, coverage plots as in Figure 1 were produced for the five F0 and F1 samples. High-coverage variants that segregated in the F1 generation were selected by hand and annotated in a variant list extracted from the raw read matrices. A custom script then used these annotated variants to rescore all samples for each of the 327 loci.

### Linkage map

The linkage map was constructed with Lep-MAP3 (v. 0.2) (Rastas 2017), after recoding the diplotypes BbHOM, HET, and BvHOM as genotypes AA, AC, and CC in the Lep-MAP3 input file. The most likely diplotype was coded as 1, and the (MLCHP) support estimates were provided for the other two diplotypes. We specified the three-generation pedigree in the input file in order to obtain a joint map across both F2 families. Lep-MAP3 was run with default parameters, except dataTolerace=0.001, distortionLod=0, grandparentPhase=1, and LodLimit=19. The most likely sex-averaged locus order in each linkage group (LG) was determined from 20 replicate runs of the OrderMarkers2 step using the Kosambi mapping function. Segregation distortion (χ^2^ estimates) per locus and family were calculated with Lep-MAP3. We applied the following significance thresholds to the χ^2^ data: (1) a Bonferroni correction, dividing α = 0.05 by the number of chromosome arms (24) in *Bombina (Morescalchi 1965; Manilo et al. 2006)*, as recommended by Fishman and McIntosh (2019) and (2) the Benjamini and Hochberg (1995) false discovery rate.

### Histology

F2 gonads with mesonephroi, fixed in Bouin’s solution, were dehydrated in an ethanol series, embedded in paraplast (Sigma), and sectioned. The 8 μm sections were stained with hematoxylin and picroaniline according to Debreuill’s trichrome procedure (Kiernan, 1990). Images were taken with a Nikon Eclipse E600 light microscope. Sex of individuals was assessed from gonad morphology (Piprek et al. 2010; Piprek 2013, see Figure S1). For some of the 162 samples, all ethanol accidentally evaporated just prior to embedding. This resulted in poor quality sections that made sex determination uncertain (*n* = 34) or impossible (*n* = 7).

### Finding the SD region

We estimate an SD bias that arises due to the nature of the crosses: In the F1s the SD haplotypes of the heterogametic parent are taxon-labeled. That is, given the direction of the F0 cross (male *B. variegata* x female *B. bombina*) and assuming an XY system, the F1 male passes the *B. variegata*-labeled Y haplotype to his sons and the *B. bombina*-labeled X haplotype to his daughters. At the SD locus, we therefore expect F2 males to be only BvHOM or HET and F2 females to be only BbHOM or HET, both in equal proportions. Further, the same pattern would be expected in a ZW system. We quantify this sex-homozygote bias with the following equation, where N[] is a count:

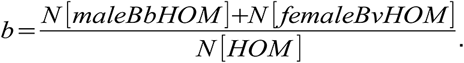

With an equal sex ratio and no heterozygote deficit, the null expectation is *b* = 0.5. At an SD (XY or ZW) locus, *b* should be zero.

In order to identify the heterogametic sex (distinguish XY from ZW systems), we needed to define a sex-limited haplotype. If this haplotype is sufficiently distinct, more than three diplotype clusters will form in the *bombina-variegata* coordinate system, with strongly sex-biased clusters. For each locus, we ranked clusters by their proportion of males, *p*_m_, and identified, in descending order, the minimal set of clusters that jointly contained more than 50% of all males. We termed the average *p*_m_ of these clusters *pMaleInMaleClusters.* At an autosomal locus, the proportion of males in each cluster will be around 0.5, and *pMaleInMaleClusters* must therefore be about 0.5. At the extreme, there may be a cluster that contains the majority of all males and no females, such that *pMaleInMaleClusters =* 1. Note that the sex-homozygote bias in the three-cluster case (BbHOM, HET, and BvHOM; see above) produces less extreme estimates. At the SD locus, the BvHOM cluster would be entirely male (*p*_m_ = 1) and contain 50% of all males. The HET cluster (expected *p*_m_ = 0.5) would need to be added to obtain more than 50% of all males, such that *pMaleInMaleClusters* would be 0.75. We similarly computed *pFemaleInFemaleClusters*.

### Data availability

Supplemental Material is currently attached to this document and will be submitted to Figshare. We will also add the complete 3-generation genotype matrix to this archive. Raw sequencing data from the WGS experiment have been submitted to ENA under study accession code PRJEB35099. Raw sequence data for all other samples and the genome assembly will be added to this.

## Results

### Genome characteristics and assemblies

From kmer frequencies (SGA (v. 0.10.15) (Simpson and Durbin 2012; Simpson 2014) preqc), we obtained a *B. variegata* genome size estimate of 7.61 Gb. A second estimate of 8.12 Gb based on the same dataset and computed with GenomeScope 2.0 (Ranallo-Benavidez et al. 2020) was provided by K.S. Jaron (pers. comm.). The average of these two, 7.87 Gb, is used throughout this paper. We explored the repeat content assembled by REPdenovo (v. 2017-02-23) (Chu et al. 2016) and extrapolated the repeats’ presence in the *B. variegata* genome based on the calculated copy number. The merged REPdenovo output contained 6,039 contigs, totaling 4.5 Mbp, with 3,689 contigs matching known Repbase repeats (Jurka et al. 2005; Bao et al. 2015). The most common repeats were *DIRS* retrotransposons (Poulter and Goodwin 2005), which were identified in 1,539 REPdenovo contigs and featured prominently in the set of 200 contigs with the highest copy number (Figure 2). The estimated total copy number of *DIRS* contigs was 807,858, covering 0.75 Gb of the *B. variegata* genome, or just under 10% of the total genome of 7.87 Gb. Other DNA transposon superfamilies that accounted for significant portions of the *B. variegata* genome included *Crypton* (0.21 Gb), *hAT* (0.19 Gb), and *Mariner* (0.10 Gb; see Table S1 for a full list). The 2,350 REPdenovo contigs that did not have any Repbase matches were estimated to cover 0.52 Gb of the *B. variegata* genome and include the REPdenovo contig with the highest copy number (Figure 2).

**Figure 2.**
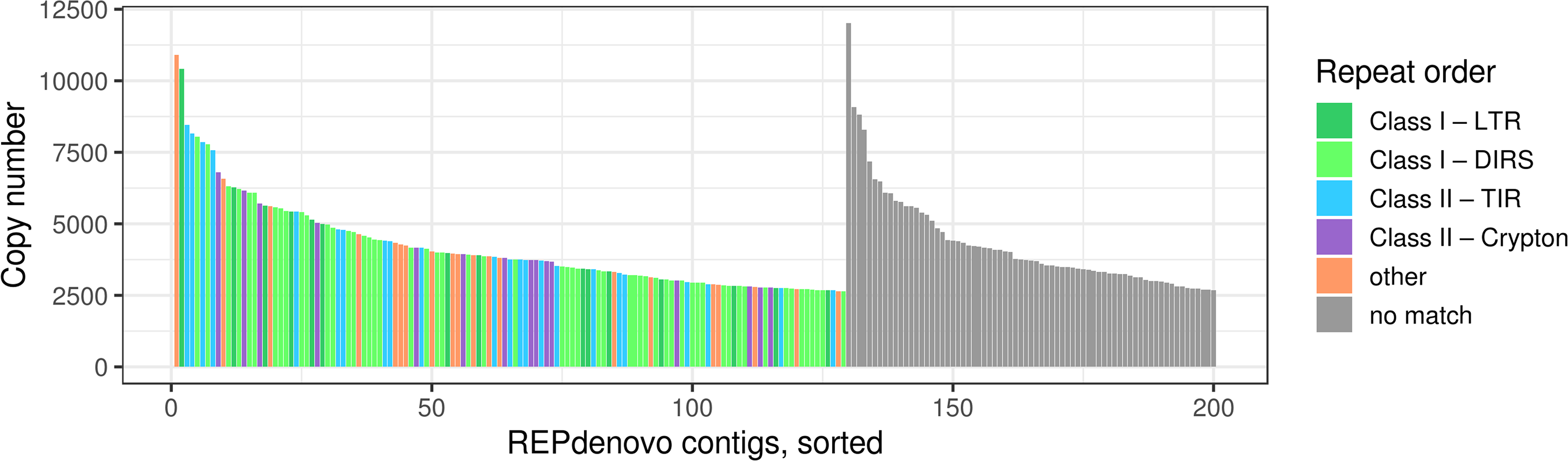
The distribution of repeat types. We show the 200 REPdenovo contigs with the highest copy number. Transposable element orders represented by more than 10 contigs in this set are identified by colour. The classification follows (Wicker et al. 2007). Contigs without a match in Repbase (blastn and tblastx) are labeled as no match and ordered separately. LTR, long terminal repeat retrotransposon; DIRS, *Dictyostelium* intermediate repeat sequence; TIR, terminal inverted repeat DNA transposon.

We assembled the *B. variegata* F0 grandfather’s genome using the CLC Genomics Workbench (v. 9.5.3) (Qiagen, Hilden, Germany), SGA (v. 0.10.15) (Simpson and Durbin 2012), and Platanus (v. 1.2.4) (Kajitani et al. 2014). CLC and SGA assembled over half of the expected genome size, though both assemblies were highly fragmented (Table 1). The Platanus assembly, which was intentionally focused on genic sequence, resulted in less than 1 Gb of contig sequence and was also extremely fragmented. Given the fragmentation, the CLC assembly was scaffolded against the *B. v. variegata* transcriptome (34,790 transcripts) with SCUBAT2 (G. Koutsovoulos, https://github.com/GDKO/SCUBAT2). SCUBAT2 assigned 73,298 CLC contigs to 13,300 paths (*i.e.* a set of contigs linked by exons from a single transcript).

**Table 1.** Assembly comparison

|  | **CLC** | **SGA** | ***Platanus*** |
| --- | --- | --- | --- |
| **Repeat-subtracted reads** | No | Yes | Yes |
| **Total contig length (Gb)** | 4.65 | 4.22 | 0.86 |
| **Number of contigs (x 10<sup>6</sup>)** | 4.37 | 7.33 | 4.59 |
| **Contig N50 length (bp)** | 1,815 | 823 | 229 |

### Reduced representation sequencing using non-repetitive baits

Candidate sequences for bait design were chosen based on uniqueness, correct assembly, and minimal redundancy, as described in the Materials and Methods. Baits were synthesised for 3,983 SCUBAT paths (including 2,407 with inferred *B. bombina* orthologues), 68 CLC contigs matching other genes of interest, and 949 CLC contigs without known gene association (total: 5,000 targets and 20,000 baits). The 4,763 ARC-assembled loci from *B. variegata*, the mapping reference, had a mean length of 673 bp, more than twice the length of the 250 bp bait region. Addition of the complete CLC contigs for the remaining 237 loci resulted in a total sequence length of 4.5 Mb.

On average, each F0, F1, or F2 sample had 1,306,372 deduplicated, on-target read pairs. Only four samples had fewer than 500,000 read pairs and belonged to one poorly performing enrichment pool. The average percentage of unique reads on target per readset was 19.8 (range: 9.5 −27.1%, excluding samples from the poorly performing pool). The average number of post-QC read pairs per sample was 4,768,367. Mapping an unenriched readset of this size to the whole genome would equate to 0.17x coverage. The observed mean coverage of the 4.5 Mb mapping reference was 147x, representing about 865-fold enrichment. The read coverage across the 5,000 targets appeared to be normally distributed (Figure S2), but we noted a potential bias when mapping the *B. bombina* grandparent reads to the separate *B. variegata* and *B. bombina* references. The average ratio of reads mapped to conspecific instead of the heterospecific reference was 1.1. However, this appeared to be the result of a small number of loci with large discrepancies (Figure S3), as the median ratio was one.

### Diplotyping and linkage mapping

Diplotypes (BbHOM, HET, BvHOM) were inferred for the two grandparents, the three F1 parents, and the 162 F2 offspring. Diplotype inference failed for 136 targets, including 77 for which no variant positions were detected. Among the 4,864 successfully clustered targets, only 25 had more than five missing diplotypes. Support estimates were greater than 10 ln likelihood units for 99.3% of the dataset (Figure S4).

Of the 4,864 targets, 4,660 were grouped into 12 LGs by Lep-MAP3, matching the published haploid chromosome number (Morescalchi 1965). We repeated the Lep-MAP3 analysis with the same dataset but replacing the data for 327 loci where the F0 grandparents and the F1 parents did not have the expected diplotype set of BvHOM, BbHOM, HET, HET, and HET. For these 327 loci, the rescored data using manually selected variants were used (see Materials and Methods). From this set, 154 were mapped in the first analysis. In the second ‘manual selection’ analysis, 138 of these 154 were placed at the same position (± 4 cM) and the remaining 16 did not map. The ‘manual selection’ analysis added 95 rescored targets to the map, bringing the total loci to 4,755 (Figure 3). This final map had a total length of 1,584 cM with 2,073 distinct map positions, separated by 0.76 cM on average.

**Figure 3.**
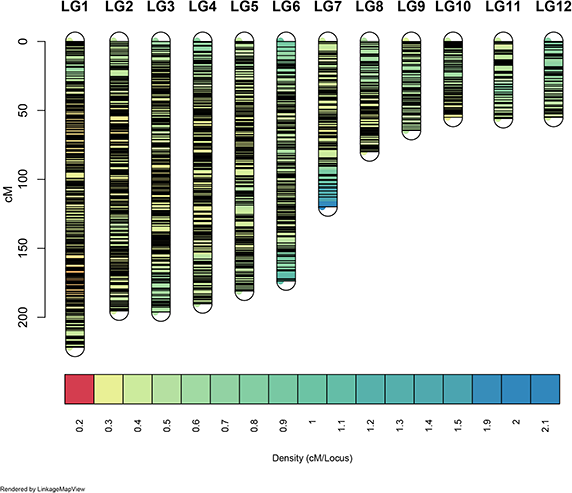
The *Bombina* linkage map. The linkage map was visualised with LinkageMapView (v. 2.1.2) (Ouellette et al. 2018). Horizontal bars represent marker loci. Colours indicate marker density in cM/locus from 0.2 (red) to 2.1 (blue).

### Segregation distortion

Across all LGs, there were eight distinct spikes in χ^2^ estimates that exceeded a lower significance threshold (the Bonferroni correction based on the number of chromosome arms), and five of these also exceeded an upper threshold (the critical value for the Benjamini and Hochberg false discovery rate; Figure 4). All eight spikes were only observed in family 6, but for some family 7 showed the same trend (LG1 right-hand spike, LG8 right-hand spike, and LG11). Based on the diplotype with the strongest deviation, there were four spikes with a HET excess, two with a BbHOM deficit and one each with a deficit and an excess of BvHOM diplotypes. Figure 4 provides χ^2^ estimates for the 4755 mapped loci, highlighting those that may be affected by scoring error.

**Figure 4.**
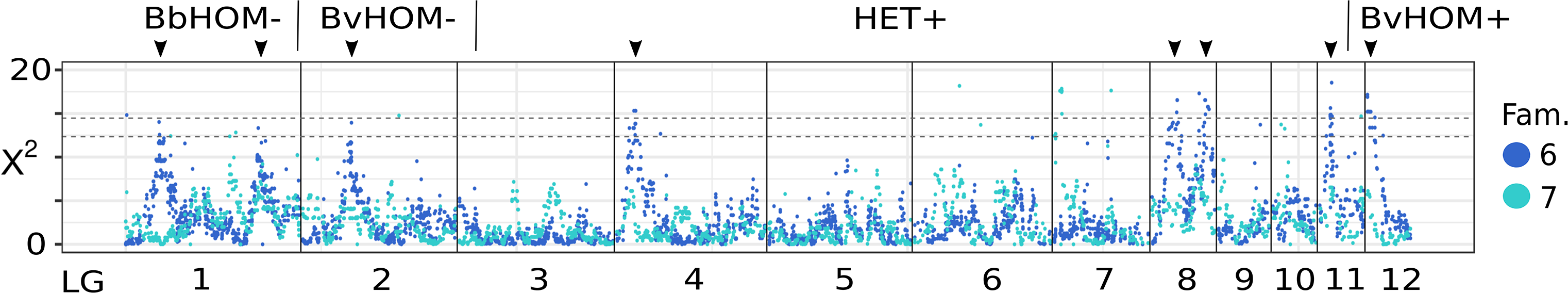
Segregation distortion, χ^2^, by family. Dashed horizontal lines are significance thresholds: the lower line is the Bonferroni correction based on the number of chromosome arms, and the upper line is the critical value for the Benjamini and Hochberg false discovery rate (the experiment-wise alpha is 0.05 in both significance thresholds). For each significant spike, which is indicated with an arrowhead, the genotype showing the strongest deviation is noted along with a (+) or (−) label, where (+) = excess and (−) = deficit. Different genotypes are separated by vertical lines above the plot. For clarity, 22 observations from 21 loci with χ^2^ > 20 are excluded from the plot.

### Large-scale synteny

We aligned the 5,000 *B. variegata* target sequences against the *X. tropicalis* genome assembly (NCBI GCA_000004195.4, Bredeson et al.) using BLAST+ (v. 2.9.0) (Camacho et al. 2009), with flags - task blastn - evalue 1E-10. Even with the large sequence divergence, 737 targets from the 12 LGs had hits to the *X. tropicalis* assembly, and the best blast hit was extracted. Although there are a small number of stray alignments, which are potentially the result of paralogy, translocations or mapping errors, the 12 LGs demonstrate obvious synteny to the *X. tropicalis* chromosomes (Figure 5). In particular, we found 1:1 correspondence between *X. tropicalis* chromosomes 1, 2, 3, 5, and 6 with LGs 2, 3, 4, 5, and 6, respectively. We also noted several distinct differences, such as intrachromosomal variation within these five conserved chromosomes or the split of *X. tropicalis* chromosome 7 into LGs 8 and 9.

**Figure 5.**
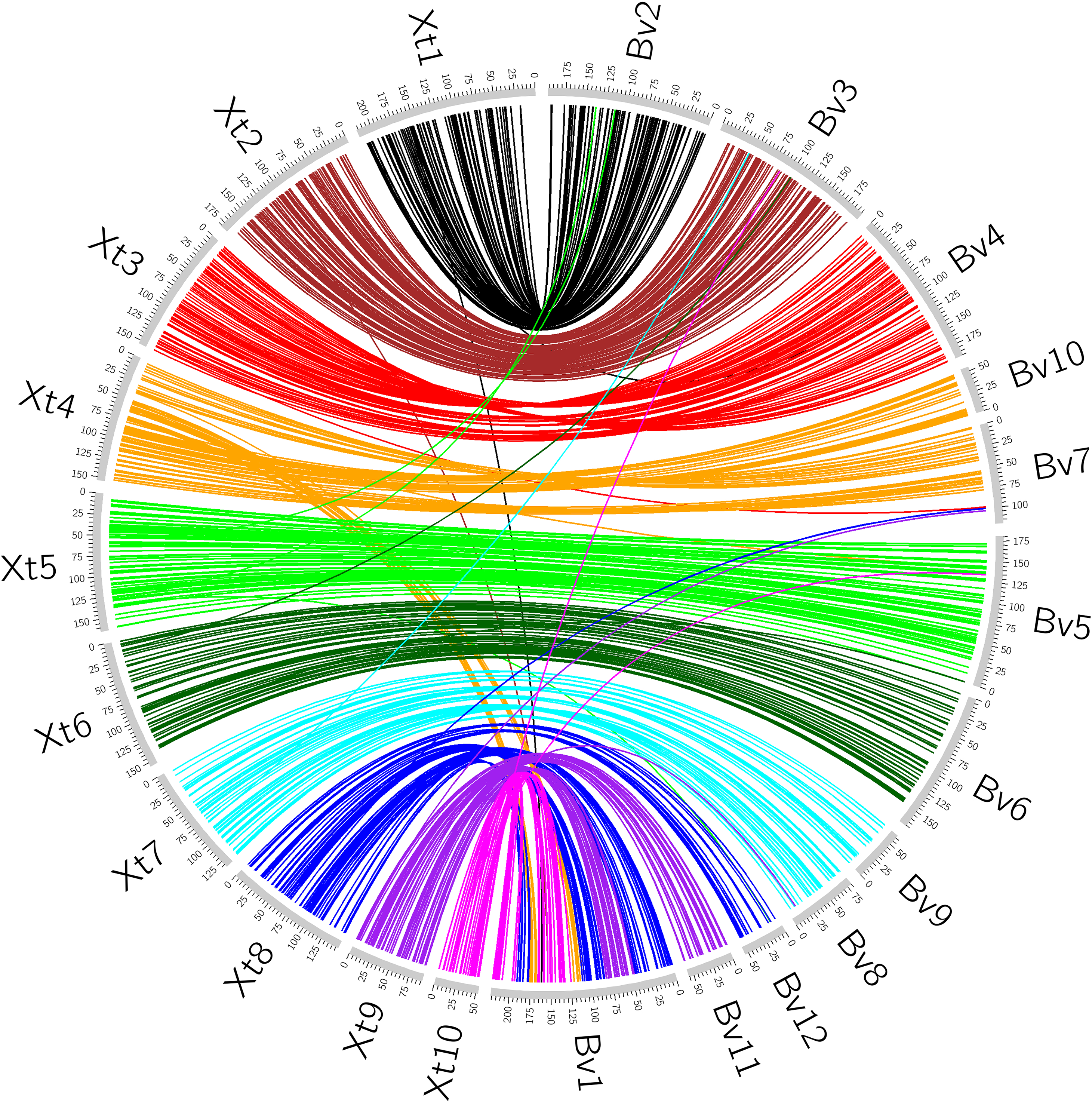
Synteny between *B. variegata and X. tropicalis*. Circos (v0.69-6) (Krzywinski et al. 2009) plot of 737 *B. variegata* target sequences from the 12 LGs (Bv, unit is cM) aligned against the *X. tropicalis* genome assembly (Xt, unit is Mb) with BLAST+ (v. 2.9.0) (Camacho et al. 2009).

### Sex-determining region

In an XY system or a ZW system, sex chromosomes would segregate in our crosses, such that males cannot be BbHOM and females cannot be BvHOM in the SD region. Therefore, we can identify the SD region based on the frequency, *b*, of these two sex-diplotype combinations among homozygotes (see Materials and Methods). The global minimum across all LGs is on LG5 at 116.09 cM (b = 0.0154), and the surrounding region (111 – 118 cM) on LG5 has a correspondingly low frequency (b < 0.017; Figure 6). Based on the null hypothesis of *b* = 0.5, this region is statistically significant with *p* < 10^−20^.

**Figure 6.**
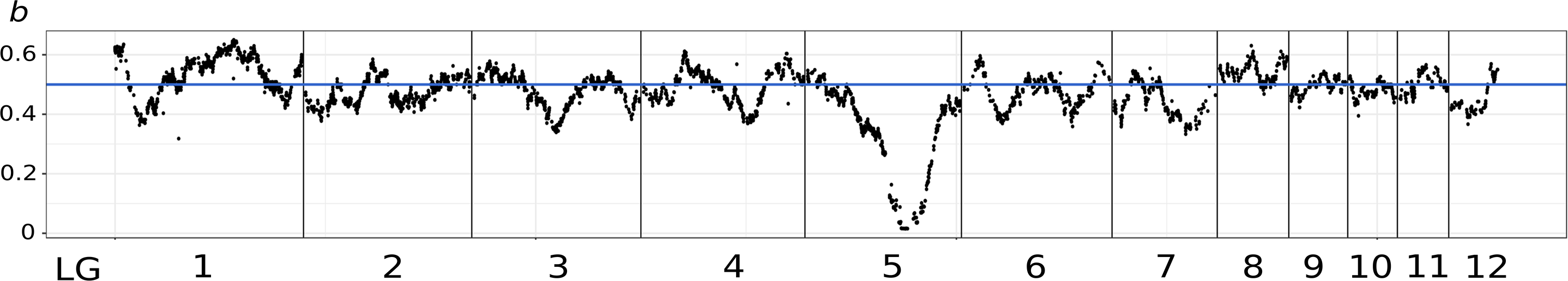
Estimated frequences of sex-diplotype combinations among homozygotes, *b*. The global minimum on LG5 indicates the sex determining region. The blue line represents the null hypothesis of b = 0.5.

In order to identify the heterogametic sex, we searched the cluster plots for instances where males were strongly associated with particular clusters, estimated as *pMaleInMaleClusters* (see Materials and Methods). This statistic had a mode at 0.5 and a mean of 0.5534. Two *pMaleInMaleClusters* outliers were identified, and both loci are located near the identified SD region. For locus 5568 (LG5, 109.33 cM), *pMaleInMaleClusters* is 0.976, and for locus 4146 (LG5, 125.00 cM), *pMaleInMaleClusters* is 0.954. We identified a strongly diverged haplotype in the male *B. variegata* grandfather at locus 5,568 (Figure 7). This haplotype was inherited by the F1 father and by 59 of the 61 F2 offspring that were unambiguously male. Only 1 of the 60 high-certainty female F2 offspring carried this haplotype. These findings imply an XY system. Closer inspection of locus 4146 revealed that the *B. bombina* grandmother had a duplication of the target region on one chromosome and a deletion on the other. This indel configuration produced the extreme *pMaleInMaleClusters* estimate (Figure S5). No outliers were observed in the analogous statistic, *pFemaleInFemaleClusters*. There is therefore no indication that *B. bombina* has a ZW system that could be competing with the *B. variegata* XY system.

**Figure 7.**
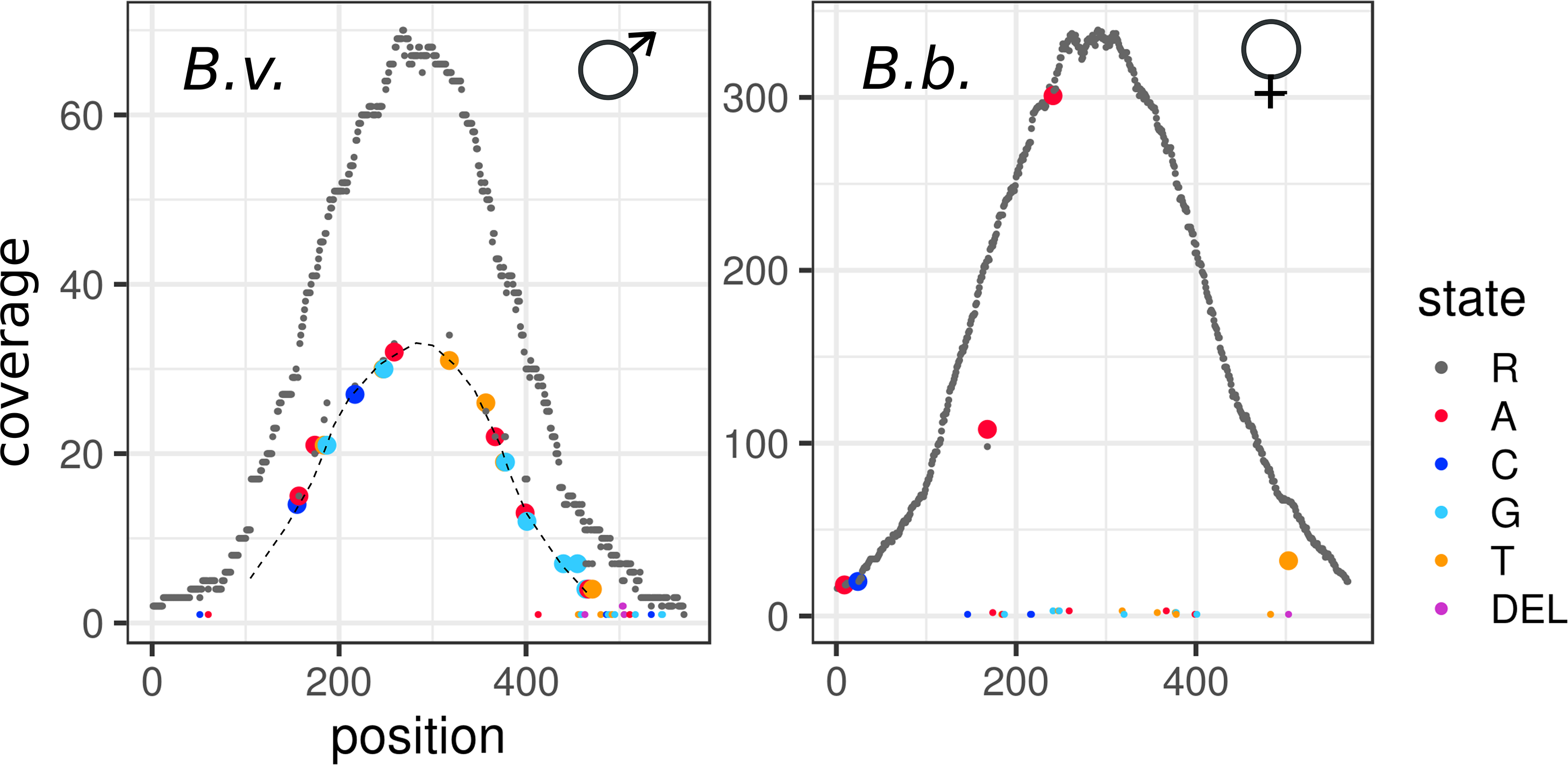
Diverged haplotype based on raw read coverage at locus 5568 in the F0 generation. Plots show the raw read coverage along the reference sequence (x-axis) for F0 *B. bombina* (left) and F0 *B. variegata* (right). Sex-linked haplotype variants in the *B. variegata* grandfather are connected with a dashed line.

## Discussion

We present here a dense *Bombina* linkage map, based on variants segregating in *B. bombina* x *B. variegata* F2 crosses. To create this linkage map, we developed a new set of molecular baits that target 4,755 loci selected from non-repetitive regions in a *de novo B. variegata* genome assembly. We inferred the most likely diplotype (BvHOM, HET or BbHOM) for each locus and sample from the raw read mapping data through a novel delayed-calling approach (cf. Nielsen et al. 2012), which eschews scoring individual variants or setting arbitrary thresholds. Using the linkage map, we identified large-scale synteny between *Bombina variegata* and *Xenopus tropicalis* as well as the location of the *Bombina* SD region and the underlying SD system.

Anuran genomes are, in general, large (average size 4.7 Gb, Gregory 2020) and have extensive repeat content (over 70% in *Oophaga pumilio* (Rogers et al. 2018) and *Leptobrachium leishanense* (Li et al. 2019b)). However, repeat composition is highly variable among anurans. While DNA transposons make up the largest fraction of repeats in *X. tropicalis* (Hellsten et al. 2010) and *L. leishanense* (Li et al. 2019), LTR retrotransposons feature prominently in *Nanorana parkeri* (Sun et al. 2015) and *O. pumilio* (Rogers et al. 2018). In *Rhinella marina (Edwards et al. 2018)* and *Vibrissaphora ailaonica* (Li et al. 2019a) around 50% of the assembled repeats are unannotated. Our high coverage short-read dataset produced a highly fragmented and partial genome assembly for the *B. variegata* grandfather of our mapping crosses. Analysis of *B. variegata* repeat content identified *DIRS* retrotransposons as the most common repeat (38% of annotated repeat content), followed by terminal inverted repeat DNA transposons (15%) and *Crypton* transposons (11%). *DIRS* and *Crypton* belong to a small subset of transposable elements that use tyrosine recombinase (YR) to integrate into the genome (Poulter and Goodwin 2005). They each account for less than 2% of the repeat content in other anuran assemblies.

These repeats hampered a previous attempt at *Bombina* marker development (Nürnberger et al. 2003), and we therefore undertook additional efforts to exclude repeats in the present study. While commercial bait design routinely masks known repeats, our bait candidates were identified from genome assemblies of a repeat-subtracted read sets, filtered based on known genes and selected transcripts, and screened with assembled REPdenovo repeats that included repeats unknown to Repbase. Screening only with known repeats could have accidentally included sequence from the REPdenovo contig with the highest copy number in the bait design, as this contig had no Repbase annotation. One measure of the success of our repeat filtering strategy is that 95% of the 5,000 enrichment targets could be integrated into the linkage map.

Because target capture was not perfect, off-target reads commonly aligned to and accumulated at one or both ends of the reference sequences. These reads introduced heterozygous variants that contradicted the variants in the centre of the reference. This was expected for a highly repetitive genome and our delayed-calling analysis pipeline was designed accordingly. Overmerging adds noise to the inheritance signal at a locus, reducing the power to call an individual’s genotype. However late-calling eschews this low power early calling step: haplotypes were instead called from the combined read data of all individuals in a homozygous cluster (~ 40), and thus at >1000-fold coverage (see Materials and Methods). When *N* is this large, the inheritance signal will dominate majority consensus calling, despite an opposing overmering signal. The converse would imply that the overmerging and inheritance signal labels are swapped. Given that baits were designed from the *B. variegata* genome assembly, we also expect enrichment bias in heterozygous individuals. With delayed-called haplotypes, we allow for such bias by maximising the likelihood of an individual’s data over the admixture coefficient between haplotype pairs, co-estimating bias. Genotype (diplotype) calls are thus late, powerful, and robust to both overmerging and enrichment bias.

While the delayed calling stage of our analyses follows standard likelihood approaches, it relies on an initial automated clustering of individual’s raw data. To asses the properties of this clustering heuristic we rescored a subset of 327 (6.5%) of loci by direct inspection, *i.e.* those that did not show the expected (BvHOM, BbHOM, HET, HET, and HET) diplotype estimates in the F0 and F1 generations (see Materials and Methods). Although such deviations are not necessarily problematic, this subset included some challenging loci. Structural variation was common, mainly homozygous or heterozygous whole-locus deletions, most of which could not be mapped. A number of loci had strongly distorted segregations and remained unmapped after rescoring. Among the loci that were added to the map (*n* = 95), there were 70 for which more than three diplotype clusters had been inferred, reflecting distinct haplotypes (alleles and/or overmergings) within one or both of the grandparents. These 70 represent about 25% of such loci on the map. While the analysis pipeline is set up to extract haplotypes from more than two clusters and compares all candidate pairs within the likelihood framework, within-taxon sequence variation appears to be the most difficult case for the clustering heuristic. This is not surprising, given its design for between-taxon variation. Nonetheless at locus 5568, the heuristic produced the same partition of the data as direct inspection, despite the strongly diverged *B. variegata* haplotype (Figures 7 and S6). Moreover, the rescoring of loci that were part of the original map brought little change: 90% of these loci were placed at essentially the same map position as before.

Overall, there were few loci with larger than expected segregation distortion (Figure 4). We report χ^2^ estimates per locus and family in Table S2 to assist future analyses. The χ^2^ spikes (Figure 4) may reflect hybrid incompatibilities or, especially in cases of homozygote deficit in one taxon, inbreeding depression in the full-sib F1 crosses (Fishman and McIntosh 2019). There were, however, no significant genotype associations between pairs of loci from different χ^2^ spikes (analyses not shown).

Our comparison between the *Bombina* linkage map and the *X. tropicalis* genome assembly provides insights into the likely Bombinanura ancestral chromosome state, and subsequent evolution, and further informs us regarding the error rate of the constructed linkage map. The observed 1:1 synteny between five *X. tropicalis* chromosomes and five *Bombina* LGs suggest that these chromosomes were present in the Bombinanura ancestor and that the distinct chromosome boundaries have been maintained for the past ~200 million years (Feng et al. 2017). The observed differences are similarly informative, suggestive of either biological diversity or linkage map construction error. If we assume the *Bombina* map estimation is error free for the five concordant chromosomes, and errors are Poisson distributed in the intervals between 732 markers, evenly distributed over 12 chromosomes, then the map error rate estimate is 0.015. This estimate is conservative, because the five ‘error free’ chromosomes have more markers than assumed. Future exploration of these synteny patterns, particularly in comparison against additional chromosome-scale frog assemblies (Mudd 2019), will increase our understanding of anuran chromosome evolution. Since frogs are a documented example of karyotypic conservatism or chromosomal bradytely (Bush et al. 1977; Baker and Bickham 1980; Marks 1983), we expected low chromosome variation between *X. tropicalis* and *Bombina*, though our visualization of these results is remarkably stark. This large-scale synteny as well as the presence of only a few stray alignments, all of which appear to be single, isolated hits, suggests that the overall structure of the linkage map agrees with the *X. tropicalis* chromosome structure and substantiates the linkage map construction.

We searched for a *Bombina* SD region using the association between homozygote genotypes and sex in F2 offspring. The same rationale was applied to recent linkage maps of *Aedes aegypti* (Fontaine et al. 2017) and *X. tropicalis* (Mitros et al. 2019). We determined the *Bombina* SD region (LG5, 111–118 cM) and at nearby locus 5568 (LG5, 109.61 cM), we identified a haplotype in the F0 *B. variegata* male that is strongly associated with male sex in the F2 generation, indicating an XY system. A preliminary analysis of *B. bombina* and *B. variegata* samples from Romania, Poland, and the Czech Republic (*n* = 35 per taxon) showed that the observed sex-linkage of this haplotype is fortuitous. In wild-caught *B. variegata*, it occured at a frequency of 0.13 and in both males and females. Male heterogamety was also established for *Bombina orientalis* (Kawamura and Nishioka 1977), the nearest relative of *B. bombina* and *B. variegata* (MRCA ~4.6 Ma; Nürnberger et al. 2016).

Similar to the situation in fish (Volff et al. 2007; Gammerdinger and Kocher 2018), the identity of the sex chromosome in amphibians can vary between closely related species and even among populations within a species (Miura 2017; Jeffries et al. 2018). Nonetheless, not all chromosomes are equally likely to take on the SD role. In anuran XY systems, chromosome 1 (numbering by homology with *X. tropicalis*) features disproportionately across diverse genera, such as *Rana, Hyla*, and *Bufo* (Brelsford et al. 2013; Tamschick et al. 2014; Miura 2017; Jeffries et al. 2018). All other known XY cases involve chromosomes 2, 3, and 5 and within the genus *Rana* switches to chromosome 5 occur more often than expected by chance (Jeffries et al. 2018). Also, known genes of the SD pathway are located on chromosome 1 (*Dmrt1, Amh*) and 5 (*FoxL2*, Jeffries et al. 2018). The observed pattern could arise if a relatively small number of genes in the vertebrate sex determination cascade alternated in assuming the master SD role (Volff et al. 2007; Graves and Peichel 2010; Herpin and Schartl 2015; Furman and Evans 2016). The *Bombina* sex chromosome is indeed homologous to *X. tropicalis* chromosome 5, but the *FoxL2* ortholog marker is located at 39.83 cM, well outside the SD region. Thus, the *Bombina* SD gene is presently unknown.

Our ability to delineate the SD region relied on the heterogametic recombination rate. In fact, the gradual decline of *b* towards its global minimum on LG5 (Figure 6) was caused entirely by recombination in the F1 male. Chiasma counts in *B. variegata* (Morescalchi 1965; Morescalchi and Galgano 1973) suggest that the female:male crossover rate is around 1.3 and that recombination in either sex is not localised to particular chromosome regions. These observations contrast with the findings in other anurans, such as *Rana, Hyla* and *Xenopus* (Brelsford et al. 2016a; b; Furman and Evans 2018), where the female recombination rate exceeds that in males up to four-fold (in one case even 75-fold, Rodrigues et al. 2013) and male crossovers are largely restricted to chromosome ends. The latter ‘recombination landscape’ is common in vertebrates (Sardell and Kirkpatrick 2020). It should favour XY sex chromosome turnover (Jeffries et al. 2018; Sardell and Kirkpatrick 2020) and contribute to the typically greater differentiation near chromosome centres relative to the ends between closely related species (Haenel et al. 2018; Sardell and Kirkpatrick 2020). We expect that these dynamics play a lesser role in *Bombina*.

The age of the *Bombina* SD system could be inferred from a phylgenetic analysis of sex linkage across sister taxa. Alternatively, X-Y sequence divergence could be estimated from loci in the non-recombining region (Charlesworth et al. 2005). However, none of the loci in the 7 cM interval where *b* is at or near its minimum had sex-linked haplotypes and are therefore presumably bracketing the SD region. Conceivably, the X and Y sequences closely associated with the SD locus are so diverged that they cannot be mapped and the non-recombining region is ‘invisible’ on the linkage map. Because there were no alignment gaps in the *X. tropicalis* chromosome 5 homologous region (Fig. 5), we suspect that this region is not very large. A small non-recombining region would be consistent with a young SD system but not proof, because some old SD systems provide counterexamples (e.g. Vicoso et al. 2013)

While whole genome sequence represents the ultimate genomic resource, it is rarely attainable and commonly non-essential. For many evolutionary questions it is sufficient to sample populations for small portions of genomes placed on a linkage map. This is particularly true for genome-wide hybrid zone studies, where linkage disequilibria require analysis in a map context but increased SNP detection provides no additional information after all segregating ancestry tracts have been marked. This applies irrespective of genome size. The approach is therefore particularly attractive for hybridising species with large genomes, provided that markers from the non-repetitive part of the genome can be identified and reliably scored. The new *Bombina* linkage map fulfills these criteria. Knowledge of the SD region and of the large-scale synteny with *X. tropicalis* broadens our scope for inference. In short, the map provides the much needed tool to take the analysis of this classic study system to a new level.

## Acknowledgements

We thank C. Chu, S, Hunter, P. Rastas and J. Simpson for advice on the use of their software. I. Jaron provided informatic support and interfacing with Czech MetaCentrum computational resources via the IVB Fishery environment. MetaCentrum Acknowledgement: Computational resources were supplied by the project “e-Infrastruktura CZ” (e-INFRA LM2018140) provided within the program Projects of Large Research, Development and Innovations Infrastructures. K.S. Jaron kindly carried out the GenomeScope analysis. A. Devault (ArborBiosciences) generously shared her bait design expertise. We thank the staff of Edinburgh Genomics for expert support in data generation. D. Podkowa prepared the histological sections and R. Piprek kindly rescored ambiguous gonad preparations in blind trials. This research used the National Energy Research Scientific Computing Center, a Department of Energy Office of Science User Facility supported by contract number DE-AC02-05CH11231. A.B.M. was supported by NIH grants R01GM086321, R01HD080708, T32GM007127, and T32HG000047 and a David L. Boren Fellowship. Collecting permits were issued by the Regional Director of Environmental Protection, Republic of Poland, OP-I.6401.193.2013.MMr. The study was approved by the First Local Ethical Committee on Animal Testing, Jagiellonian University, Kraków (94/V/2013 nr 86/2013). We acknowledge financial support from the Polish National Science Centre (grants 2013/09/B/NZ8/03349; UMO-2013/09/B/NZ8/03349) to J.M.S. and B.N. and from the Czech Science Foundation (grant 16-26714S) to B.N.

